# Radical amino acid mutations persist longer in the absence of sex

**DOI:** 10.1101/049924

**Authors:** Joel Sharbrough, Meagan Luse, Jeffrey L. Boore, John M. Logsdon, Maurine Neiman

## Abstract

Harmful mutations are ubiquitous and inevitable, and the rate at which these mutations are removed from populations is a critical determinant of evolutionary fate. Closely related sexual and asexual taxa provide a particularly powerful setting to study deleterious mutation elimination because sexual reproduction should facilitate mutational clearance by reducing selective interference between sites and by allowing the production of offspring with different mutational complements than their parents. Here, we compared the rate of removal of conservative (*i.e*., similar biochemical properties) and radical (*i.e*., distinct biochemical properties) nonsynonymous mutations from mitochondrial genomes of sexual *vs*. asexual *Potamopyrgus antipodarum*, a New Zealand freshwater snail characterized by coexisting and ecologically similar sexual and asexual lineages. Our analyses revealed that radical nonsynonymous mutations are cleared at higher rates than conservative changes and that sexual lineages eliminate radical changes more rapidly than asexual counterparts. These results are consistent with reduced efficacy of purifying selection in asexual lineages allowing harmful mutations to remain polymorphic longer than in sexual lineages. Together, these data illuminate some of the population-level processes contributing to mitochondrial mutation accumulation and suggest that mutation accumulation could influence the outcome of competition between sexual and asexual lineages.

## INTRODUCTION

One of the primary hypothesized advantages of sexual reproduction is the clearance of harmful mutations, which is expected to be much more effective when linkage disequilibria (LD) are disrupted by sex (Hill and Robertson 1966). Although this LD disruption should be most apparent in the nuclear genome, biparental inheritance and meiotic recombination in the nuclear genome should also decrease LD between the nuclear genome and the mitochondrial genome (*i.e*., mitonuclear LD) because nuclear and mitochondrial genomes can be transmitted separately. One important consequence of the potential for separate transmission of nuclear and mitochondrial alleles is a substantial reduction in selective inference between the mitochondrial and nuclear genome relative to a scenario (*e.g*., most forms of asexual reproduction) where these genomes are always or nearly always co-transmitted (Normark and Moran 2000; Neiman and Taylor 2009). The evolutionary consequences of sexual reproduction (here, increased efficacy of natural selection) should therefore extend beyond the nuclear genome to cytoplasmic genomes, which are typically uniparentally transmitted and often lack recombination (Barr et al. 2005). Elevated mitonuclear LD resulting from co-transmission of nuclear and mitochondrial genomes, combined with simultaneous selection on nuclear variants, should impede selection in the cytoplasmic genomes of asexual lineages (Normark and Moran 2000).

Reduced efficacy of selection in the cytoplasmic genomes of asexual lineages is most obviously linked to accumulation of *slightly* deleterious mutations over time (Gabriel et al. 1993; Neiman and Taylor 2009); however, the Hill-Robertson effect (*i.e*., smaller *N_e_*s*) is expected to reduce the efficacy of selection with respect to *all* mutational changes. The implications are that although most strongly deleterious mutations should eventually be purged from asexual populations, these mutations will tend to remain polymorphic for a longer period of time than similar mutations in sexual populations. Thus even severely deleterious changes will have a higher probability of fixation by random drift in asexual *vs*. sexual lineages.

The prediction that asexual lineages should experience a higher rate of accumulation of harmful mutations has found support from animal (Neiman et al. 2010; Henry et al. 2012) and plant (Voigt-Zielinski et al. 2012; Hollister et al. 2015; Lovell et al. 2017) taxa. While these results represent important steps towards understanding the genomic consequences of asexuality, the evolutionary mechanisms (*e.g*., deleterious mutation accumulation *vs*. mitonuclear coevolution) underlying these observations remain unclear, in large part because the extent to which accumulated mutations are actually deleterious in asexuals has not been evaluated. This type of information is especially important in light of substantial evidence that nonsynonymous mutations are likely to vary widely in fitness effects (Eyre-Walker and Keightley 2007; Boyko et al. 2008). Predicting and characterizing the fitness effects of these nonsynonymous mutations remains a central area of focus in evolutionary biology (Keightley and Charlesworth 2005; Xue et al. 2008; Eyre-Walker and Keightley 2009; Andolfatto et al. 2011; Halligan et al. 2011). Here, we take advantage of the fact that the biochemical properties of amino acids represent useful heuristics (*e.g*., conservative *vs*. radical amino acid changes) for inferring effects of mutations on protein phenotype (Zhang 2000; Hanada et al. 2007; Popadin et al. 2007) to take critical steps towards assessing whether mutations likely to influence fitness accumulate differently in asexual than in sexual lineages.

The removal of deleterious mutations (and the fixation of beneficial mutations) depends upon the efficacy of selection as well as the fitness effect of mutations, such that elevated *N_e_* and reduced LD in sexual *vs*. asexual populations should result in more rapid removal of deleterious mutations for the former (Birky and Walsh 1988). As such, comparing rates and patterns of evolution across reproductive modes while incorporating information about mutational severity will provide a powerful glimpse into the evolutionary dynamics governing the removal of deleterious mutations. The New Zealand freshwater snail *Potamopyrgus antipodarum* is ideally suited to evaluate this critically important evolutionary process because otherwise similar obligately sexual and obligately asexual *P. antipodarum* frequently coexist within New Zealand lake populations (Lively 1987; Jokela et al. 1997), enabling direct comparisons across reproductive modes, and, thereby, across genomes that experience predictable variation in the efficacy of selection.

Asexual lineages of *P. antipodarum* are the product of multiple distinct transitions from sexual *P. antipodarum* (Neiman and Lively 2004; Neiman et al. 2011), meaning that these asexual lineages represent separate natural experiments into the consequences of the absence of sex. Neiman *et al*. (2010) showed that asexual lineages of *P. antipodarum* experience a higher rate of nonsynonymous substitution in their mitochondrial genomes than sexual lineages. Here, we extend these studies to a relatively short evolutionary time scale by using whole mitochondrial genomes to evaluate whether sexual lineages distinguish between radical (*i.e*., ancestral and derived amino acids have distinct biochemical properties) and conservative (*i.e*., ancestral and derived amino acids have similar biochemical properties) amino acid changes more effectively than asexual lineages. This approach allowed us to evaluate whether harmful mutation accumulation is detectable within species and, if so, whether this phenomenon is driven by ineffective selection in asexual lineages. The outcome of these analyses emphasizes fundamental differences in the rate of accumulation of conservative *vs*. radical nonsynonymous mutations and the critical importance of intraspecific sampling. Our results also demonstrate that radical mutations persist longer in mitochondrial genomes of asexual lineages of *P. antipodarum* compared to sexual counterparts, likely as a consequence of reduced efficacy of purifying selection.

## MATERIALS & METHODS

### Sequencing

We analyzed 31 whole mitochondrial genomes from eight sexual lineages and 23 asexual lineages of *P. antipodarum* and one whole mitochondrial genome from the closely related *Potamopyrgus estuarinus*, representing the wide-ranging mitochondrial genetic diversity of this species in New Zealand (see Figure 1, Table S1; Neiman and Lively 2004; Neiman et al. 2010; Paczesniak et al. 2013). We obtained 19 publicly available mitochondrial genomes (4 sexual, 14 asexual, 1 *P. estuarinus*, Accession Nos.: GQ996415 – GQ996433, Neiman et al. 2010), sequenced eight mitochondrial genomes (2 sexual, 6 asexual) using bi-directional Sanger sequencing on an ABI 3730 (Applied Biosystems, Foster City, CA), and sequenced and assembled five mitochondrial genomes (2 sexual, 3 asexual) using 2x100 bp paired-end sequencing on an Illumina HiSeq 2500 (Illumina, Inc., San Diego, CA) as part of the ongoing *P. antipodarum* nuclear genome project. For all newly sequenced lineages, we determined ploidy and, thus reproductive mode (2x – sexual; ≥ 3x – asexual), using flow cytometry following the protocol outlined in (Neiman et al. 2011; Neiman et al. 2012; Paczesniak et al. 2013; Krist et al. 2014) and extracted total genomic DNA following a mollusk-adapted phenol-chloroform extraction protocol (Fukami et al. 2004). We cleaned DNA extractions using the Zymo Clean and Concentrate Kit (Zymo Research, Irvine, CA), re-suspended DNA in 30-100 µL of T-low-E buffer (10mM Tris pH8.0, 0.1mM EDTA), and determined DNA concentration and purity for each sample on a NanoDrop 1000 (Thermo Fisher Scientific, Waltham, MA). For the eight Sanger-sequenced samples, we amplified mitochondrial genomes in four overlapping fragments using primers and programs designed in Neiman et al (2010). PCR products were cleaned with Shrimp Exo shrimp alkaline phosphatase (Werle et al. 1994) and directly sequenced with internal sequencing primers (Table S2). Sanger-sequenced mitochondrial genomes were assembled and manually edited in Sequencher 5.0 (Gene Codes Corporation, Ann Arbor, MI). Only unambiguous sites supported by ≥ 2 reads were included in the assemblies.

**Figure 1.**
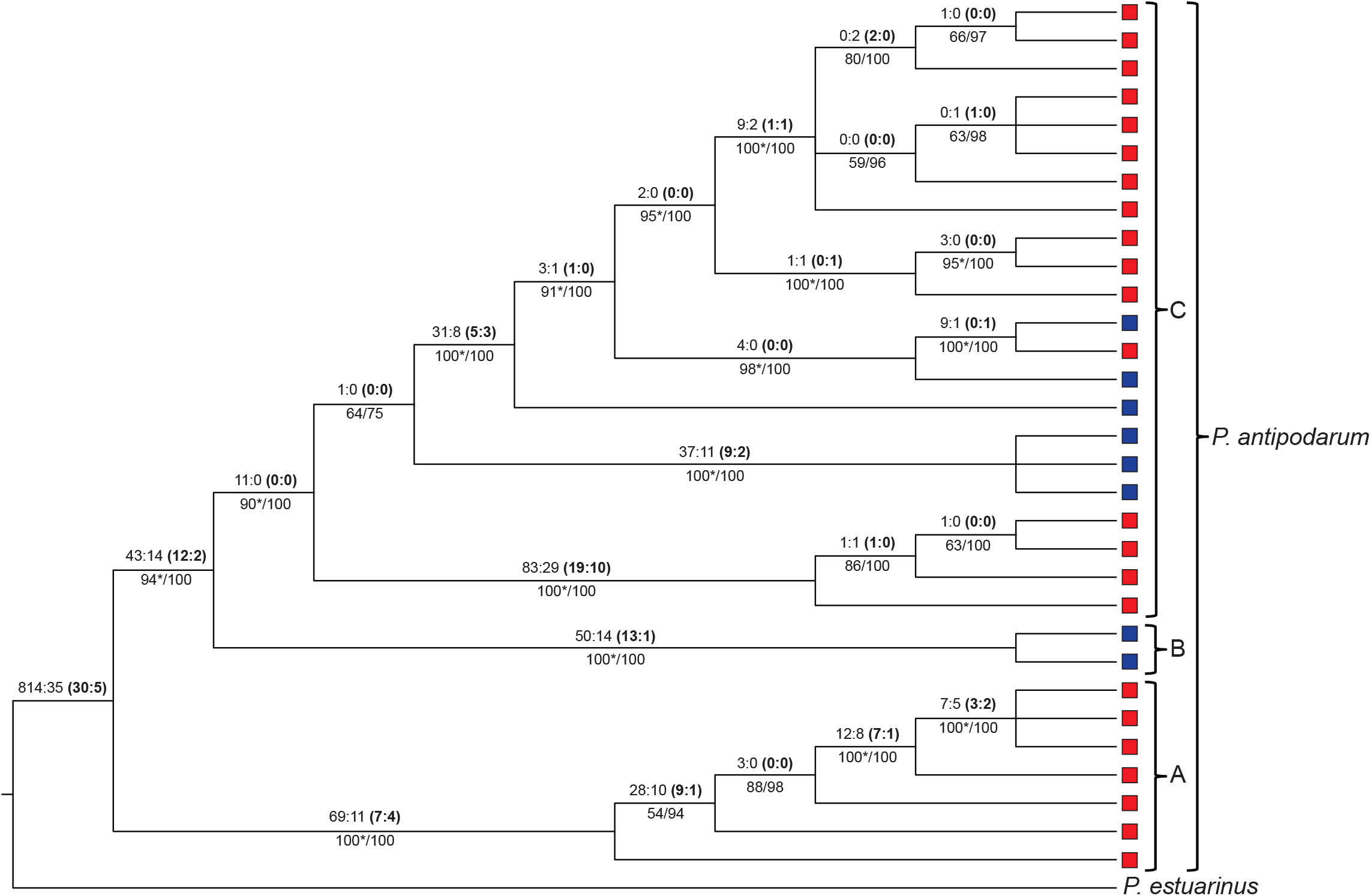
Whole-mitochondrial genome phylogeny for *P. antipodarum* and an outgroup, *P. estuarinus*. Consensus phylogenetic tree depicting evolutionary relationships of 23 asexual (red) and eight sexual (blue) *P. antipodarum* mitochondrial genomes. Tree topology was inferred using Maximum Likelihood (ML, performed in PAUP*) and Bayesian (performed in MrBayes) methods. Two models of molecular evolution identified by jModelTest v 2.0 (GTR + I + G, TIM3 + I + G) were used to infer ML-based trees, while the model of molecular evolution was directly inferred from the alignment over 1.0 × 10^6^ generations for the Bayesian (MrBayes) tree. Branch support from the GTR + I + G ML approach and from the Bayesian consensus tree is listed below branches (bootstrap support/posterior probability). Asterisks indicate nodes for which both ML models exhibited ≥ 90 bootstrap support. Numbers above each line represent the numbers of non-homoplasious mutational changes, with the ratio of synonymous: nonsynonymous changes in plain font and the ratio of conservative: radical changes in bold font and parentheses. Clade A represents a particularly distinct group of asexual lineages.

For the remaining five DNA samples, we constructed sequencing libraries using the Illumina Nextera DNA Library Prep Kit and sequenced 2x100 bp paired-end reads on a single lane of an Illumina HiSeq 2500. Following read trimming and quality control (≥ 30 Phred quality score) using the Fastx-toolkit (Gordon and Hannon 2010), we assembled mitochondrial genomes *de novo* using the CLC Genomics Work Bench (Qiagen, Hilden, Germany). Consensus base calls were then determined for sites with >10x coverage and Phred-scaled quality scores >30. Sites with >1 read supporting a minor allele (resulting from *e.g*., heteroplasmies or *numts*) were not used in this analysis. All newly sequenced mitochondrial genomes have been deposited in GenBank (Accession Nos.: *sequences will be uploaded to GenBank upon acceptance*).

### Phylogenetic analysis

We aligned whole mitochondrial genome sequences from the 31 *P. antipodarum* lineages and one *P. estuarinus* sample using MAFFTv v7.305b (Katoh and Standley 2013) and manually edited the alignments in MEGA v5.2.2 (Kumar et al. 2008, Supplementary File S2). We used jModelTest 2, which uses a corrected Akaike Information Criterion (AICc), Bayesian Information Criterion (BIC), and performance-based decision theory (DT) in a Maximum Likelihood (ML) framework, (Darriba et al. 2012), to select appropriate models of molecular evolution. Of the 1,624 possible models evaluated by jModelTest 2, two models produced the highest log-likelihood scores across all three model estimators: TIM3 (Posada 2003) and GTR (Tavaré 1986), both with invariant sites (I) and a gamma distribution of site classes (G). We therefore used both models (TIM3+I+G, GTR+I+G) to infer ML-based phylogenies using PAUP*4.0 (Swofford and Sullivan 2009) with 1,000 bootstrap replicates each and assuming the following priors: TIM3+I+G: lset – base = (0.2587, 0.1736, 0.1748), nst = 6, rmat = (2.7926, 49.1045, 1.0000, 2.7926, 37.8686), rates = gamma, shape = 0.8150, ncat = 4, pinvar = 0.4550; GTR+I+G: lset – base = (0.2597, 0.1736, 0.1738), nst = 6, rmat = (1.5661, 37.1103, 0.5913, 2.9024, 28.5161), rates = gamma, shape = 0.8380, ncat = 4, pinvar = 0.4610. We also inferred the phylogeny for these mitochondrial genomes under a Bayesian statistical framework using MrBayes v3.2 (Ronquist et al. 2012) assuming the following priors: nucmodel = 4by4, nst = mixed, ploidy = haploid, rates = invgamma, ngammacat = 4, number of generations = 10^6^, relative burn-in = 25%, number of chains = 4, heating parameter = 0.5, revmat = all GTR submodels have equal probabilities, statefreq = dirichlet, shape = exponential (1.0), pinvar = uniform (0.0, 1.0), ratemultiplier = fixed (1.0), topology = all topologies equally probable, brlens = unconstrained: gammaDir (1.0, 0.1, 1.0, 1.0). After ensuring that the number of generations (10^6^ generations) and burn-in (2.5 × 10^5^ generations) were sufficient for the log probability of trees to plateau, trees were sampled every 500 generations, resulting in 24,016 trees of which 2,045 contributed to the 95% credible tree set. We visualized ML majority rule trees and the consensus Bayesian tree in FigTree v1.4 (Rambaut 2012); only nodes with bootstrap values ≥ 60 and posterior probabilities ≥ 75 were used in subsequent tests of molecular evolution (Figure 1). Although animal mitochondrial genomes are small (~15 kbp), there can still be substantial among-gene heterogeneity in evolutionary rate (Xia 1998; Ballard 2000; Marshall et al. 2009), which can affect phylogenetic inferences (Maddison 1997; Mallo et al. 2016). We therefore used MrBayes v3.2 to infer gene trees for each of the 13 protein-coding genes, each with their own empirically determined substitution model, in the *P. antipodarum* mt genome. We then evaluated whether nodes not present in the genome-wide tree had significant statistical support (posterior probabilities ≥ 75) in individual gene trees.

Recombination can make phylogenetic analysis complex or even impossible (Schierup and Hein 2000; Arenas and Posada 2010a, b; Arenas and Posada 2014; Mallo et al. 2016), even in mitochondrial genomes (Rokas et al. 2003; Barr et al. 2005; Tsaousis et al. 2005). Recombination in mitochondrial genomes is particularly problematic in phylogenetic analyses of sexual lineages *vs*. asexual lineages because only sexual lineages have the potential to experience paternal leakage and recombination between mitochondrial genomes from different lineages. To evaluate evidence for recombination among *P. antipodarum* mitochondrial genomes, we estimated population recombination rate using LDHat (implemented by RDP4 – Martin et al. 2017). To test whether recombination had any effect on our phylogenetic analyses, we also inferred a genome-wide phylogenetic tree using MrBayes excluding those regions with evidence of recombination.

Next, we extracted and concatenated the 13 protein-coding regions from the alignment (~11.2 kbp, 3,738 amino acids) for all 32 lineages, and we mapped all mutational changes in protein-coding regions to the fixed tree topology (described above) according to the rules of parsimony. Only non-homoplasious changes were considered in subsequent phylogenetic analyses of molecular evolution. In our population genetic analyses, we considered all sites that were variable within *P. antipodarum* (including homoplasious sites) to be polymorphic and all sites that were distinct between *P. antipodarum* and *P. estuarinus* to be substitutions.

### Identifying conservative and radical amino acid changes

We used seven different amino acid classification schemes, three drawn from Zhang (2000), three drawn from Hanada *et al*. (2007), and a modified Grantham scheme based on charge and hydrophobicity (Grantham 1974), to evaluate rates and patterns of radical and conservative amino acid evolution (Table 1). While there is some overlap between different classification schemes, each scheme highlights different amino acid properties that are likely to shape protein evolution. For example, amino acid charge is a major determinant of protein folding (Perutz et al. 1965; Anfinsen 1973; Nakashima et al. 1986; Bashford et al. 1987; Wright et al. 2005) and three-dimensional structure (Lesk and Chothia 1980; Geisler and Weber 1982; Doms et al. 1988; Rumbley et al. 2001), changes between polar and non-polar amino acids can expose or bury key interaction residues in membrane-associated proteins (von Heijne 1992), and volume and aromaticity can both affect protein folding and play a role in protein-protein interactions (Burley and Petsko 1985). Classification schemes 4 and 7 (Table 1) are unique in that they are based on evolutionary information (although classification scheme 7 largely fits with charge and polarity classifications), meaning that these schemes incorporate aspects of other amino acid characteristics into their classifications. Using these seven amino acid classification schemes, we developed an overall index of the degree of amino acid change that we termed the Conservative-Radical Index (CRI). For each possible amino acid change, we calculated CRI by averaging across the seven amino acid classification schemes (radical changes assigned a value = 1.0, conservative changes assigned a value = 0.0), such that CRI = 1.0 indicates that the amino acid change in question was radical in all amino acid classification schemes (*e.g*., V ← → E) whereas CRI = 0.0 indicates that the amino acid change in question was conservative in all amino acid classification schemes (*e.g*., V ← → I). Amino acid changes with CRI ≤ 0.5 were treated as conservative changes and amino acid changes with CRI > 0.5 were treated as radical changes (Table S3). All analyses of molecular evolution and population genetics were repeated for each amino acid scheme individually as well as for the classification scheme-averaged classifier.

**Table 1.** Amino acid classification schemes.

| 1 | 2 | 3 | 4 | 5 | 6 | 7 |
| --- | --- | --- | --- | --- | --- | --- |
| Zhang<br>(2000) | Zhang<br>(2000) | Zhang<br>(2000) | Hanada et<br>al. (2007) | Hanada et<br>al. (2007) | Hanada et<br>al. (2007) | Grantham<br>(1974) |
| Charge | Polarity | Polarity &<br>Volume | Correlation<br>with $K_A/K_S$ | Charge &<br>Aromaticity | Charge &<br>Polarity | Charge &<br>Hydrophobicity |
| *Neutral*<br>A, C, F,<br>G, I, L, M,<br>N, P, Q,<br>S, T, V,<br>W, Y | *Polar*<br>C, D, E, G,<br>H, K, N, Q,<br>R, S, T, Y | *Neutral, small*<br>A, G, P, S,<br>T | *Neutral, small*<br>A, C, G, N,<br>P, S, T | *Neutral, non-aromatic*<br>A, C, G, I, L,<br>M, N, P, Q,<br>S, T, V | *No polarity*<br>A, F, G, I,<br>L, M, P,<br>V, W | *Neutral, hydrophobic*<br>A, F, I, L, M,<br>P, V, W |
| *Positively charged*<br>H, K, R |  | *Polar, small*<br>D, E, N, Q | *Basic, acid, aromatic, small*<br>F, H, K, Q,<br>R, W, Y | *Neutral, aromatic*<br>F, W, Y | *Neutral, polar*<br>C, N, Q,<br>S, T, Y | *Neutral, hydrophilic*<br>C, G, N, Q,<br>S, T, Y |
| *Negatively charged*<br>D, E | *Non-polar*<br>A, F, I, L,<br>M,<br>P, V, W | *Nonpolar, small*<br>I, L, M, V | *Neutral, large*<br>I, L, M, V | *Basic*<br>H, K, R | *Basic, polar*<br>H, K, R | *Positive, hydrophilic*<br>H, K, R |
|  |  | *Polar, large*<br>H, K, R |  |  |  |  |
|  |  | *Nonpolar, large*<br>F, W, Y | *Acidic, large*<br>D, E | *Acidic*<br>D, E | *Acidic, polar*<br>D, E | *Negative, hydrophilic*<br>D, E |
|  |  | *Special*<br>C |  |  |  |  |

We then calculated the number of mutational target sites per codon for each different type of mutational change (*i.e*., synonymous, nonsynonymous, conservative nonsynonymous, and radical nonsynonymous) relative to the invertebrate mitochondrial genetic code (http://www.ncbi.nlm.nih.gov/Taxonomy/Utils/wprintgc.cgi#SG5) to ensure that we properly accounted for the different probabilities of different types of mutational changes (Table S4). To confirm that the number of each of type of site was properly calculated, we checked that the number of nonsynonymous sites and the number of synonymous sites per codon summed to three and that the number of conservative nonsynonymous sites and the number of radical nonsynonymous sites per codon summed to the number of nonsynonymous sites per codon. Of particular note is that the “GTG”, “TTG”, “ATT”, “ATC”, and “ATA” codons can all be used as alternative start codons in invertebrate mitochondrial genomes (only the GTG alternative start codon was observed in the present dataset), which we accounted for in our site calculations.

### Molecular evolution and population genetic analyses

To evaluate the relative intensity of selection acting on different types of mutational changes, we estimated rates of substitution between *P. antipodarum* and *P. estuarinus* (Li et al. 1985), ratios of polymorphism to divergence (McDonald and Kreitman 1991), nucleotide diversity (Watterson 1975; Nei and Li 1979; Fu and Li 1993), and site frequency spectra (SFS, Tajima 1989) in protein-coding regions of the *P. antipodarum* mitochondrial genome for each mutational type using all seven amino acid classification schemes and the scheme-averaged classifier with a custom python tool (available at https://github.com/jsharbrough/conservativeRadicalPolymorphismsSubstitutions). For codons with multiple hits, we assumed the fewest number of nonsynonymous changes necessary to explain the data.

To compare rates of molecular evolution in *P. antipodarum* relative to *P. estuarinus*, we first calculated substitution rates for each mutational type in *P. antipodarum* as in Li *et al*. (1985), corrected these estimates for multiple hits using the Jukes-Cantor model of molecular evolution (Jukes and Cantor 1969), and estimated variance as in Jukes and Cantor (1969). We then compared 95% confidence intervals (CIs) of substitution rates between *P. antipodarum* and *P. estuarinus*, estimated as 95% 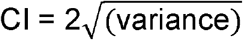 for each substitution type, such that non-overlapping CIs indicate statistically different rates of substitution. We also took advantage of the non-homoplasious changes mapped to our fixed tree topology (see above) to compare synonymous substitution rate (*K_S_*), nonsynonymous substitution rate (*K_A_*), conservative substitution rate (*K_C_*), and radical substitution rate (*K_R_*) relative to *P. estuarinus*. We estimated branch lengths by summing the mapped changes for each lineage and mutational type and dividing the number of changes by the number of mutational target sites (Table S4). We then used pairwise Wilcoxon Signed-Rank (WSR) tests and the Holm procedure for correcting for multiple comparisons to determine whether branch lengths differed across mutational types (Holm 1979). We next used McDonald-Kreitman (MK) tests to compare molecular evolution in conservative *vs*. radical changes using a series of Fisher’s Exact Tests (FET): (1) synonymous changes to conservative changes, (2) synonymous changes to radical changes, and (3) conservative changes to radical changes. All statistical tests were performed in R v3.2.4 (R Core Team 2016).

To compare species-wide patterns of polymorphism, we estimated nucleotide diversity using *θ*_*π*_ – hereafter *π* – (Nei and Li 1979) and *θ_W_* – hereafter *θ* – (Watterson 1975), and their respective variances (Durrett 2008) within *P. antipodarum*. We compared levels of nucleotide diversity across mutational types by constructing 95% CIs for each mutational type using 10,000 bootstrap replicates. For each bootstrap replicate, we randomly selected 3,738 codons with replacement and recalculated all population genetic statistics for each replicate using a custom Python tool (available at https://github.com/jsharbrough/popGenBootstrapping). We also performed an estimate of *θ* using only private polymorphisms – hereafter *θ_U_*. Because private polymorphisms should often represent relatively young mutations that are relatively independent of phylogenetic constraint (Fu and Li 1993), these polymorphisms can be used to estimate rates of mutation accumulation since the transition to asexuality. We compared levels of these private polymorphisms across mutational types with the same bootstrapping approach as with *π* and *θ*, using a custom Python tool to compare CIs for each mutational type and compute *p* values (available at https://github.com/jsharbrough/distributionPvalues). This tool rank-orders bootstrap replicates and tests the degree to which bootstrap distributions overlap (*i.e*., perfect overlap of bootstrap distributions: *p >* 0.9998, no overlap of bootstrap distributions: *p* < 0.0002, for a two-tailed test). Because deleterious changes should generally be found at lower frequencies than relatively neutral changes, we also compared the SFS across mutational types using a series of Goodness-of-Fit (GoF) analyses assuming *X^2^* distribution, first for nonsynonymous, conservative, and radical polymorphisms *vs*. synonymous polymorphisms, and second for conservative *vs*. radical polymorphisms. We performed these GoF analyses with the chisq function in R v3.2.4 (R Core Team 2016) and visualized species-wide polymorphism levels using the R package ggplot2 (Wickham 2016).

After establishing that the substitution rate for radical amino acid changes is lower than for conservative amino acid changes and that radical changes exhibit lower nucleotide diversity within species than conservative changes (see Results, Figure 2), we compared rates of mutation accumulation of conservative and radical amino acid changes across reproductive modes. We used the mapped changes to estimate synonymous-corrected branch lengths for conservative and radical sites and compared rates of molecular evolution for each mutational type separately in sexual *vs*. asexual lineages using Mann-Whitney U (MWU) tests. We also performed these same comparisons on internal branches only and on branch tips only to test whether patterns of mutation accumulation across reproductive modes were robust across evolutionary time.

**Figure 2.**
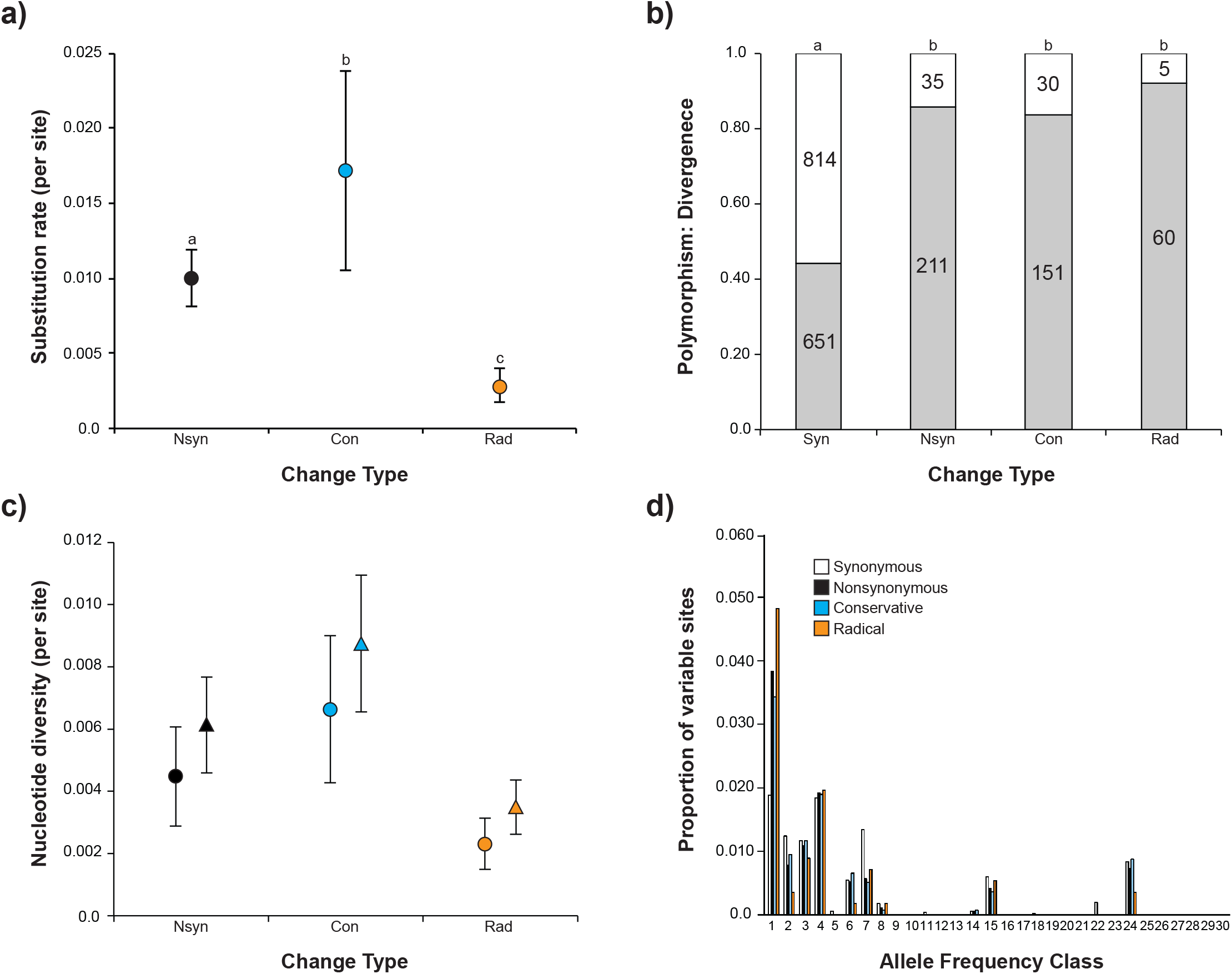
Molecular evolution and population genetics of synonymous, nonsynonymous, conservative, and radical mutations in *P. antipodarum* mitochondrial genomes. a) Branch-specific estimates of synonymous-corrected substitution rates for all nonsynonymous changes (black), conservative nonsynonymous changes (blue), and radical nonsynonymous changes (orange), as identified by the model-averaged amino acid classification scheme (CRI). Error bars represent standard deviations of branch-specific substitution rates. Lower-case letters reflect statistical groupings based on pairwise Mann-Whitney U tests (*p* < 0.017). b) Ratios of polymorphism to divergence for synonymous, all nonsynonymous, conservative nonsynonymous, and radical nonsynonymous changes. Lower-case letters reflect statistical groupings based on pairwise Fisher’s Exact Tests (*p* < 0.013). c) Nucleotide diversity estimated using *π* (circles) and *θ* (triangles) for all nonsynonymous (black), conservative nonsynonymous (blue), and radical nonsynonymous (orange) sites. Error bars reflect variance calculated as in Durrett (2008). Radical nonsynonymous sites exhibit significantly lower levels of nucleotide diversity than conservative sites (*p* < 0.0002) for both measures of polymorphism. d) Site frequency spectra of synonymous (white), nonsynonymous (black), conservative nonsynonymous (blue), and radical nonsynonymous (orange) polymorphisms within *P. antipodarum*.

We next compared polymorphism levels for conservative and radical changes across reproductive modes by dividing our sample into sexual (n = 8) and asexual (n = 23) groups and separately calculating *π*, *θ*, and *θ_U_* for each mutational type. Asexual *P. antipodarum* have already been shown to accumulate nonsynonymous changes more rapidly than their sexual counterparts (Neiman et al. 2010), leading to the a priori expectation that asexual lineages should exhibit higher levels of nucleotide diversity at nonsynonymous sites than sexual lineages. To address whether this prediction was met, we calculated a test statistic, *D_AS_*, in which 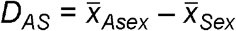, where *x̄* represents the diversity-related population genetic statistic under consideration. We then constructed a null distribution for each of these statistics using a custom-built python program (available at https://github.com/jsharbrough/conRadNullDistribution), which randomly assigned lineages without replacement and without consideration of reproductive mode to one of two groups: group 1 (n = 23, representing the asexual lineage sample size) and group 2 (n = 8, representing the sexual lineage sample size), calculated the difference between the two groups, *D_12_*, where 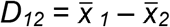, for each population genetic statistic, and repeated this process 10,000 times. Because we expected asexual lineages to exhibit higher levels of nonsynonymous polymorphism than sexual lineages, we conducted one-tailed tests of polymorphism across reproductive modes (H_O_: *D_AS_* = 0; H_A_: *D_AS_* > 0) by comparing *D_AS_* to the null distribution for each population genetic statistic (*i.e.*, *π*, *θ*, and *θ_U_*) for synonymous sites, synonymous-corrected conservative sites, and synonymous-corrected radical sites. We inferred *p* values directly from these comparisons using a custom python tool (available at https://github.com/jsharbrough/distributionPValues) and visualized null distributions and test statistics using the ggplot2 package for R (Wickham 2016).

## RESULTS

### Phylogenetic analysis of mitochondrial genetic diversity in *P. antipodarum.*

We used 32 whole mitochondrial genomes to infer a phylogeny of sexual and asexual *P. antipodarum* relative to a *P. estuarinus* outgroup using Maximum Likelihood and Bayesian methods (Figure 1). Because we did not detect any major differences from our genome-wide tree in individual gene trees (Figure S1) or in a tree that excluded regions suggestive of recombination (Figure S2), we performed our molecular evolutionary analyses assuming the genome-wide tree topology.

These phylogenetic analyses revealed substantial intraspecific genetic variation within *P. antipodarum* (mean pairwise genetic distance ≈ 0.025 changes/site) as well as relatively deep divergence for one entirely asexual clade (clade A; mean pairwise distance between clade A *vs*. clades B/C ≈ 0.035 changes/site, *vs*. mean pairwise distance within B/C ≈ 0.013). Clade A is composed mostly of lineages from the North Island of New Zealand (Lakes Waikaremoana and Tarawera) but also includes one invasive lineage from Wales and two lineages collected from lakes on the South Island of New Zealand (Lakes Brunner and Kaniere). Clades B/C are predominantly composed of lineages collected from the South Island of New Zealand (Lakes Alexandrina, Brunner, Grasmere, Gunn, Heron, Ianthe, Kaniere, Lady, McGregor, Poerua, Rotoiti, Rotoroa); however, two invasive lineages (collected from Lake Superior in Duluth, MN, USA and Denmark), one North Island lineage (collected from Lake Waikaremoana), and a lineage collected from Lake Alexandrina form a monophyletic, all-asexual clade within clade C.

We next used parsimony to map the 1,524 non-homoplasious changes (out of 1,711 total variable sites) present in the ~11 kbp protein-coding region of the mitochondrial genome onto this tree topology (Figure 1). In all, there were 814 synonymous substitutions and 35 nonsynonymous substitutions between *P. antipodarum* and *P. estuarinus*. Of these 35 nonsynonymous substitutions, 30 were considered conservative amino acid changes and 5 were considered radical amino acid changes by our scheme-averaged classifier index, CRI (see Materials and Methods). Within *P. antipodarum*, we observed 651 synonymous polymorphisms and 211 nonsynonymous polymorphisms; CRI classified 151 of the latter as conservative amino acid polymorphisms and the other 60 nonsynonymous polymorphisms as radical amino acid changes (Figure 2b).

### Substitution rates and polymorphism levels for conservative *vs*. radical amino acid changes

Using the mapped changes to calculate branch-specific estimates of Jukes-Cantor-corrected substitution rate (*K*), we found that mean branch-specific synonymous substitution rate (*K_S_*) was significantly higher than mean branch-specific nonsynonymous substitution rate (*K_A_*; WSR: *V* = 496, *p* < 0.0001), conservative substitution rate (*K_C_*; WSR: *V* = 496, *p* < 0.0001), and radical substitution rate (*K_R_*; WSR: *V* = 496, *p* < 0.0001), revealing that nonsynonymous changes evolve at lower rates than do synonymous changes (Table 2). Branch-specific estimates of *K_C_* were also significantly higher than branch-specific estimates of *K_R_* (WSR: *V* = 496, *p* < 0.0001), indicating that among amino acid-changing mutations, radical amino acid changes evolve particularly slowly (Figure 2a, Table 2). In accordance with branch-specific estimates, species-wide estimates of substitution rates that only use fixed differences between *P. antipodarum* and *P. estuarinus* revealed that *K_S_* was significantly greater than *K_A_*, *K_C_*, and *K_R_*, and that *K_C_* was significantly greater than *K_R_* (Table S5). In summary, branch-specific and species-wide estimates of substitution rate indicate that nonsynonymous sites have lower rates of substitution than synonymous sites and that radical nonsynonymous sites have lower rates of substitution than conservative nonsynonymous sites.

**Table 2.** Substitution rate and nucleotide diversity across mutational types in *P. antipodarum* mitochondrial genomes. For each mutational type, we estimated branch-specific substitution rates (*K*), nucleotide diversity (*π* and *θ*), and private allele diversity (*θ_U_*) using seven distinct amino acid classification schemes plus a model-averaged classifier (CRI).

| Mutational Type | $K$<br>(Variance<sup>a</sup>) | $\pi^b$<br>(95% CI) | $\theta^b$<br>(95% CI) | $\theta_U^b$<br>(95% CI) |
| --- | --- | --- | --- | --- |
| Syn | 0.48 *<br>(0.46 – 0.51) | 0.064 *<br>(0.058 – 0.069) | 0.063 *<br>(0.058 – 0.067) | 0.0087 *<br>(0.0069 – 0.011) |
| Nsyn | 0.0077<br>(0.0060 – 0.0093) | 0.0046<br>(0.0038 – 0.0053) | 0.0061<br>(0.0052 – 0.0069) | 0.0025<br>(0.0020 – 0.0031) |
| Con<sup>c</sup> | 0.013<br>(0.010 – 0.015) | 0.0065<br>(0.0050 – 0.0075) | 0.0083<br>(0.0067 – 0.0094) | 0.0034<br>(0.0022 – 0.0036) |
| Rad<sup>c</sup> | 0.0059 **<br>(0.0034 – 0.0085) | 0.0025 ***<br>(0.0018 – 0.0034) | 0.0037 ***<br>(0.0028 – 0.0047) | 0.0017<br>(0.00076 – 0.0024) |
| Con-1 | 0.0087<br>(0.0072 – 0.010) | 0.0058<br>(0.0047 – 0.0066) | 0.0074<br>(0.0062 – 0.0084) | 0.0029<br>(0.0019 – 0.0034) |
| Rad-1 | 0.0049 **<br>(0.0017 – 0.0081) | 0.0015 ***<br>(0.00078 – 0.0024) | 0.0026 ***<br>(0.0016 – 0.0037) | 0.0015<br>(0.0016 – 0.0033) |
| Con-2 | 0.0094<br>(0.0078 – 0.011) | 0.0055<br>(0.0040 – 0.0061) | 0.0065<br>(0.0053 – 0.0075) | 0.0026<br>(0.0022 – 0.0042) |
| Rad-2 | 0.0052 **<br>(0.0029 – 0.0076) | 0.0037<br>(0.0028 – 0.0048) | 0.0055<br>(0.0044 – 0.0068) | 0.0024<br>(0.0016 – 0.0028) |
| Con-3 | 0.015<br>(0.012 – 0.017) | 0.0077<br>(0.0060 – 0.0094) | 0.0094<br>(0.0076 – 0.011) | 0.0032<br>(0.0018 – 0.0036) |
| Rad-3 | 0.0042 **<br>(0.0026 – 0.0057) | 0.0030 ***<br>(0.0022 – 0.0036) | 0.0045 ***<br>(0.0035 – 0.0052) | 0.0022<br>(0.0017 – 0.0031) |
| Con-4 | 0.0145<br>(0.011 – 0.018) | 0.0069<br>(0.0054 – 0.0084) | 0.0084<br>(0.0069 – 0.010) | 0.0027<br>(0.0025 – 0.0042) |
| Rad-4 | 0.0032 **<br>(0.0022 – 0.0043) | 0.0031 ***<br>(0.0022 – 0.0037) | 0.0046 ***<br>(0.0036 – 0.0054) | 0.0024 <sup>e</sup><br>(0.0011 – 0.0024) |
| Con-5 | 0.011<br>(0.0095 – 0.013) | 0.0075<br>(0.0061 – 0.0088) | 0.0093<br>(0.0078 – 0.011) | 0.0033<br>(0.0020 – 0.0037) |
| Rad-5 | 0.0041 **<br>(0.0017 – 0.0064) | 0.0017 ***<br>(0.00095 – 0.0022) | 0.0028 ***<br>(0.0020 – 0.0035) | 0.0017<br>(0.0016 – 0.0029) |
| Con-6 | 0.012<br>(0.0094 – 0.014) | 0.0055<br>(0.0040 – 0.0064) | 0.0067<br>(0.0053 – 0.0078) | 0.0029<br>(0.0020 – 0.0038) |
| Rad-6 | 0.0041 **<br>(0.0025 – 0.0058) | 0.0039<br>(0.0030 – 0.0048) | 0.0056<br>(0.0046 – 0.0067) | 0.0022<br>(0.0016 – 0.0030) |
| Con-7 | 0.014<br>(0.011 – 0.016) | 0.0062<br>(0.0046 – 0.0073) | 0.0075<br>(0.0059 – 0.0087) | 0.0029<br>(0.0024 – 0.0040) |
| Rad-7 | 0.0029 **<br>(0.0017 – 0.0041) | 0.0034 ***<br>(0.0026 – 0.0043) | 0.0050<br>(0.0041 – 0.0060) | 0.0023<br>(0.0012 – 0.0025) |

<sup>a</sup> – Variance calculated as standard deviation multiplied by a factor of two
<sup>b</sup> – 95% CIs estimated from 10,000 bootstrap replicates
<sup>c</sup> – Mutational type defined by CRI score
\* – All nonsynonymous mutational types had significantly lower values ( $p < 0.05$ after correcting for multiple comparisons) than synonymous sites for all estimates of polymorphism and substitution
\*\* – Radical substitution rate was significantly lower ( $p < 0.0001$ ) than conservative substitution rate
\*\*\* – Nucleotide diversity was significantly lower ( $p < 0.05$ after correcting for multiple comparisons) at radical sites than at conservative sites

To determine the probability of fixation of conservative *vs*. radical nonsynonymous polymorphisms, we performed pairwise MK tests of selection across all mutational types. We found that nonsynonymous (FET: *p* < 0.0001), conservative (FET: *p* < 0.0001), and radical (FET: *p* < 0.0001) amino acid polymorphisms were significantly less likely to reach fixation than synonymous polymorphisms (Figure 2b, Table 3). There was a trend for conservative polymorphisms to become fixed within *P. antipodarum* at higher rates than radical polymorphisms, but this difference was not significant (FET: *p* = 0.098). These data indicate that nonsynonymous changes have a lower probability of fixation than synonymous changes, but that there is no difference in fixation probabilities across different types of nonsynonymous changes.

**Table 3.** McDonald-Kreitman tests of selection across mutational types in *P. antipodarum* mitochondrial genomes.

|  | Syn | Nsyn (Nsyn/Syn) | Con<sup>a</sup> (Con/Syn) | Rad<sup>a</sup> (Rad/Syn) |
| --- | --- | --- | --- | --- |
| Polymorphism | 651 | 211 (0.32) | 151 (0.23) | 60 (0.092) |
| Divergence | 814 | 35 (0.043) | 30 (0.037) | 5 (0.0061) |
| FET vs. Syn | - | $p < 0.0001$ | $p < 0.0001$ | $p < 0.0001$ |
| FET vs. Con | - | $p = 0.59$ | - | $p = 0.098$ |

<sup>a</sup> – Mutational type defined by CRI score

To investigate how conservative and radical changes evolve within species, we estimated nucleotide diversity for all mutational types using the frequency-dependent measure *π*, the frequency-independent measure *θ*, and a measure of private nucleotide diversity *θ_U_*, treating all lineages as a single population. Comparisons of *π* revealed that nonsynonymous, conservative, and radical sites exhibited lower levels of nucleotide diversity than synonymous sites (*p* < 0.0002 for all comparisons, Figure S3a, Table 2). Similarly, nonsynonymous, conservative, and radical sites all exhibited lower levels of nucleotide diversity than synonymous sites, as measured by *θ* (*p* < 0.0002 for all comparisons, Figure S3b, Table 2), and *θ_U_* (*p* < 0.0002 for all comparisons, Table 2, Figure S3c, Table 2). Together, these results indicate that all types of nonsynonymous changes are eliminated from *P. antipodarum* mitochondrial genomes more rapidly than synonymous changes (Figure 2c, Table 2). We also found that *π_C_* and *θ_C_* were significantly greater than *π_R_* (*p* = 0.0004) and *θ_R_* (*p* < 0.0002), respectively (Table 2, Figure 2c); however, *θ_U-C_* and *θ_U-R_* are statistically indistinguishable within *P. antipodarum* (*p* = 0.072, Figure S3c, Table 2). Because asexual lineages likely experience distinct demographic forces that sexual lineages do not (Charlesworth and Wright 2001; Kaiser and Charlesworth 2009), we also compared these population genetic statistics using only sexual lineages and found evidence of similarly elevated intensity of selection against radical *vs*. conservative polymorphisms as observed in analyses incorporating all *P. antipodarum* mitochondrial genome sequences (Figure S4).

Finally, we compared allele frequencies of different mutational types by comparing their SFS (Figure 2d). We found that nonsynonymous (*X^2^* = 56.14, df = 7, *p* < 0.0001), conservative (*χ^2^* = 28.57, df = 7, *p* < 0.0001), and radical SFS (*X^2^* = 35.04, df = 7, *p* < 0.0001) are significantly left-skewed (*i.e*., excess of rare variants) relative to synonymous changes. We did not detect any differences in conservative *vs*. radical SFS (*X^2^* = 6.36, df = 4, *p* = 0.17). In sum, all types of nonsynonymous changes are found in lower numbers and at lower frequencies than synonymous changes and that radical amino acid polymorphisms tend to be found in lower numbers and at lower frequencies than conservative amino acid polymorphisms.

### Efficacy of purifying selection in sexual *vs*. asexual lineages

To evaluate the degree to which the efficacy of selection is reduced in asexual *vs*. sexual lineages of *P. antipodarum*, we used the same set of 1,524 synapomorphic and autapomorphic changes (Figure 1) to estimate sexual and asexual lineage branch lengths relative to outgroup *P. estuarinus* and compare relative rates of conservative and radical amino acid molecular evolution. After accounting for the number of mutational target sites (Table S4) and correcting for multiple hits (Jukes and Cantor 1969), we found that asexual lineages exhibited higher rates of substitution at synonymous sites than sexual lineages (MWU: *W* = 142, *p* = 0.025, Figure S5a, Table 4), potentially indicating that the mitochondrial genomes of asexual lineages of *P. antipodarum* experience higher mutation rates than their sexual counterparts. We therefore corrected nonsynonymous rates of molecular evolution with synonymous rates to ensure that we could adequately compare efficacy of selection acting on conservative and radical changes across reproductive modes. We did not detect a difference in the rate of molecular evolution at conservative sites in sexual *vs*. asexual lineages (MWU: *W* = 95, *p* = 0.91, Figure 3a, Table 4). By contrast, we did find that asexual lineages exhibit significantly higher rates of molecular evolution at radical sites than sexual lineages (MWU: *W* = 168, *p* < 0.0001, Figure 3b, Table 4), indicating that asexual lineages accumulate radical amino acid changes more rapidly than sexual lineages. We obtained a similar pattern of accelerated accumulation of radical amino acid changes in asexual *vs*. sexual lineages when excluding branch tips from the analysis (*K_C_*/*K_S_* – MWU: *W* = 84, *p* = 0.74; *K_R_*/*K_S_* – MWU: *W* = 168, *p* < 0.0001), indicating that this pattern is robust across evolutionary time. Results from all amino acid classification schemes are detailed in Table S6.

**Table 4.**
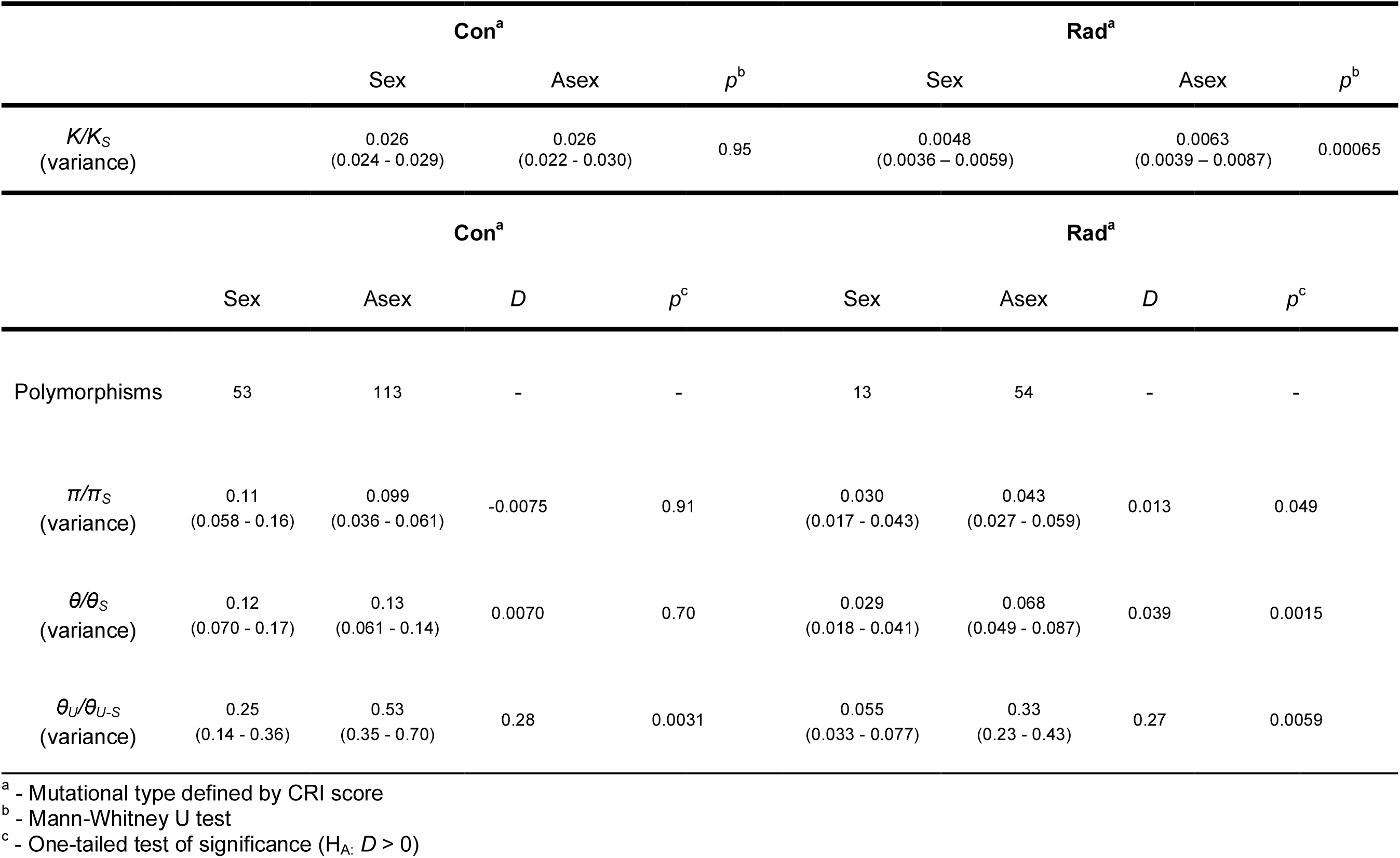
Population genetic comparisons of conservative and radical mutations in sexual vs. asexual lineages of *P. antipodarum*.

**Figure 3.**
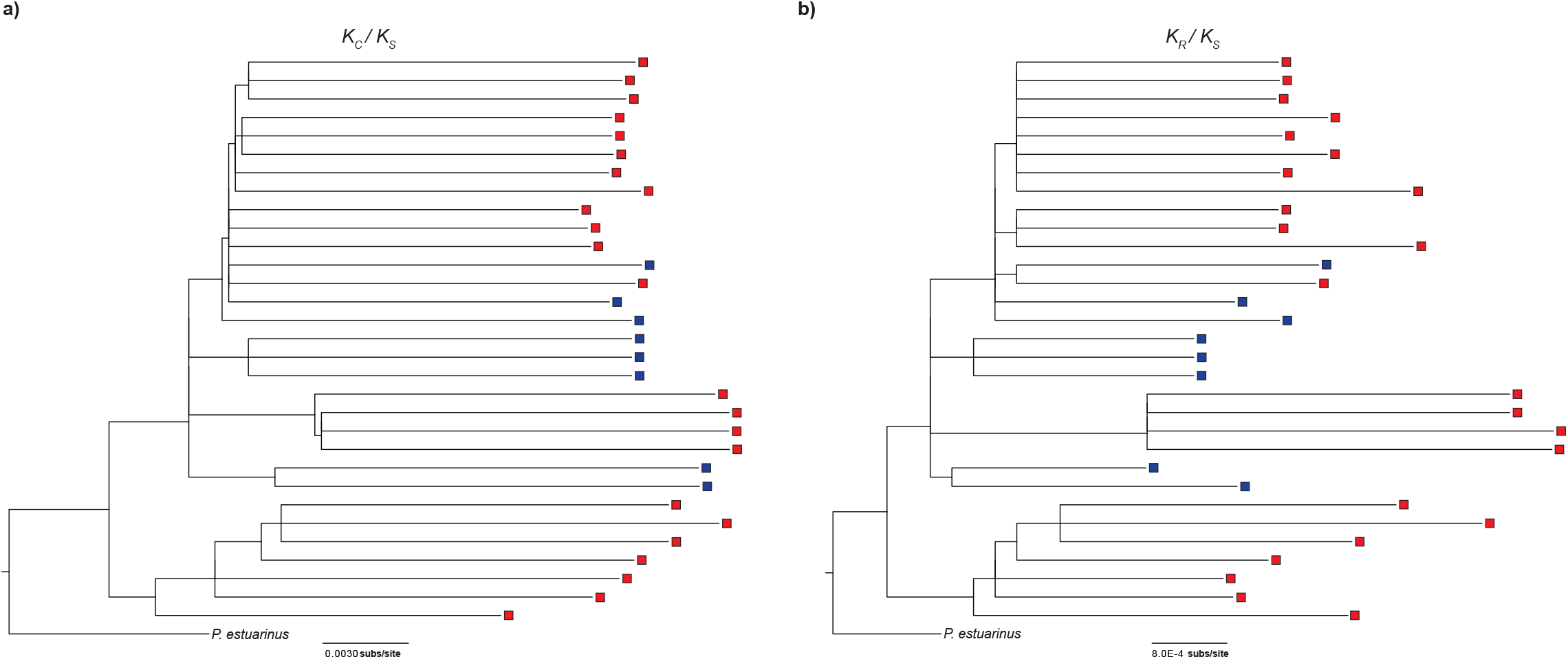
Molecular evolution of conservative and radical amino acid substitutions in *P. antipodarum*. Branch-specific estimates of a) synonymous-corrected conservative nonsynonymous substitution rates did not differ across sexual (blue) and asexual (red) lineages (*W* = 95, *p* = 0.91), but b) synonymous-corrected radical nonsynonymous substitution rates were significantly higher in asexual lineages than in sexual lineages (*W* = 168, *p* = 0.00065).

We next compared patterns of polymorphism across reproductive modes and mutational types using *π* and *θ*. For each mutational type, we calculated an estimate of nucleotide diversity for asexuals and sexuals separately, using the difference D as a test statistic to compare against a null distribution (see Materials and Methods). Although sexual and asexual lineages do not appear to differ in terms of synonymous polymorphism levels (Figure S5b, Table 4), *K_S_*, was elevated in asexual compared to sexual lineages (Figure S5a, Table 4). Because this result could reflect a scenario where differential effects of neutral processes change in response to the transition to asexuality (Charlesworth and Wright 2001; Kaiser and Charlesworth 2009), we accounted for levels of neutral polymorphism in both sexuals and asexuals by correcting nonsynonymous estimates of nucleotide diversity with the corresponding synonymous rate. For both *π* and *θ*, there was no evidence that sexual lineages differed from asexual lineages in the level of conservative amino acid polymorphism (*D_π_* = -0.0086, *p* = 0.91, Figure S6a; *D_θ_* = 0.014, *p* = 0.70, Figure 4a, Figure S6c, Table S6). The same analyses applied to radical amino acid polymorphisms revealed significantly higher levels of radical polymorphisms in asexual *vs*. sexual lineages (*D_π_* = 0.012, *p* = 0.049, Figure 4a Figure S6b; *D_θ_* = 0.035, *p* = 0.0015, Figure S6d, Table S6).

**Figure 4.**
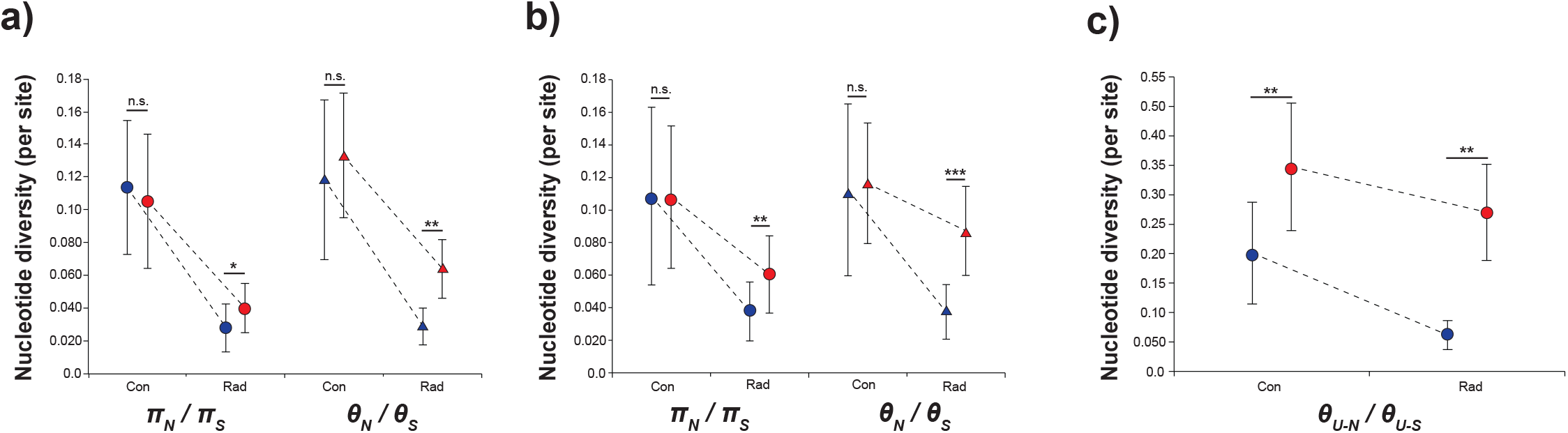
Population genetic comparisons of conservative and radical amino acid polymorphism in sexual (blue) *vs*. asexual (red) lineages of *P. antipodarum*. a) *π_N_*/*π_S_* (circles) and *θ_N_*/*θ_S_* (triangles) in sexual *vs*. asexual lineages for conservative and radical sites. b) *π_N_*/*π_S_* (circles) and *θ_N_*/*θ_S_* (triangles) in sexual *vs*. asexual lineages for conservative and radical sites. Only lineages from clade C were used in this analysis. c) Private polymorphism (*θ_U-N_/θ_U-S_*) in sexual *vs*. asexual lineages for conservative and radical sites. Error bars reflect variance calculated as in Durrett (2008). For all comparisons, one-tailed *p* values (*i.e*., Asex > Sex) are indicated by asterisks (* – *p* < 0.05, ** – *p* < 0.01, *** – *p* < 0.001).

Clade A asexuals appear to be genetically distinct relative to the rest of the *P. antipodarum* dataset (Figure 1), raising the question of whether this group might be contributing disproportionately to our observations of elevated radical amino acid polymorphism levels in asexual *P. antipodarum*. We therefore repeated the comparisons of *π* and *θ* with clade A excluded, using the clade B sexuals as an outgroup to clade C (Figure 4b, Table 4). In this more limited analysis, we again found that asexuals harbored higher levels of radical (*D_π_* = 0.023, *p* = 0.0038; *D_θ_* = 0.050, *p* < 0.0001) but not conservative amino acid polymorphism (*D_π_* = -0.00057, *p* = 0.60,; *D_θ_* = 0.0040, *p* = 0.74) (Figure 4b, Figure S6e-h), indicating that the clade A asexuals were not entirely responsible for the patterns of elevated radical amino acid polymorphism in *P. antipodarum* asexuals.

To account for the phylogenetic non-independence of asexual lineages in *P. antipodarum* in our population genetic analyses, we also compared levels of private polymorphism across reproductive modes and mutational types (Figure 4c, Table 4). There was a trend towards higher levels of synonymous private polymorphism in sexual compared to asexual lineages (*D_θU_* = -0.0017, *p* = 0.057, Figure S5b), so we corrected for private synonymous polymorphism levels before comparing private nonsynonymous sites between sexuals and asexuals. These analyses revealed that asexual lineages exhibited significantly higher levels of both private conservative amino acid polymorphism (*D_θU_* = 0.28, *p* = 0.0031) and private radical amino acid polymorphism than sexual lineages (*D_θU_* = 0.27, *p* = 0.0059, Figure 4c, Table S6). The lower levels of private polymorphism in sexual compared to asexual lineages for both conservative and radical mutational types indicates that selection is less effective in asexual lineages and that this effect is detectable even over relatively short time scales.

## DISCUSSION

### Radical amino acid changes accumulate in the absence of sex

Here, we provide evidence from a non-model system featuring reproductive mode polymorphism (a primary determinant of *N_e_*) that there exists more stringent purifying selection on radical *vs*. conservative mutational types. This pattern is pronounced enough to be evident even among intraspecific comparisons. These results are consistent with the expectation that radical mutations are usually more harmful than conservative mutations (Freudenberg-Hua et al. 2003; Smith 2003), and demonstrate that radical and conservative mutations appear to experience very different histories of selection in natural populations. The implication is that radical amino acid changes usually impart substantially more severe fitness effects than conservative changes and should therefore return to mutation-selection-drift equilibrium more rapidly than conservative amino acid changes (Fisher 1930).

We also found that asexual *P. antipodarum* lineages experience higher rates of radical amino acid substitution and exhibit higher incidences (*θ*) and frequencies (*π*) of radical amino acid polymorphisms than sexual counterparts. These results suggest that asexual *P. antipodarum* experience a relatively high rate of accumulation of likely harmful mutations in their mitochondrial genomes. Although using biochemical properties to predict likely phenotypic consequences of amino acid changes is ultimately limited by the reality that amino acid position can also play a role in protein properties (see *e.g*., Zhai et al. 2009), compensatory coevolution between nuclear and mitochondrial genomes has been documented on a background of conservative, but not radical, structural evolution (Havird et al. 2015). The implications of this latter finding are that radical amino acid changes are rarely, if ever, tolerated in the multi-subunit complexes of OXPHOS, making adaptive evolution an unlikely explanation underlying the elevated numbers of radical mutations we have observed in mitochondrial genomes of asexual *P. antipodarum*. We therefore interpret this result as evidence that mutations in asexual lineages experience lower *N_e_^*^s* than in sexual lineages, with the implication that the elimination of even strongly deleterious mutations should occur more slowly in asexual than in sexual lineages. Theoretical (Ohta 1987; Charlesworth et al. 1993; Charlesworth and Wright 2001) and empirical (Eyre-Walker and Keightley 2007; Wright et al. 2008; Katju et al. 2015) work support this conclusion, suggesting that populations with low *N_e_* should harbor a larger proportion of “effectively neutral” mutations than populations with large *N_e_*. The somewhat surprising upshot of this prediction is that even severely deleterious mutations become more effectively neutral and thus more likely to be subject to fixation by drift when *N_e_* is low. The absence of any detectable differences in sexual and asexual lineages in their rate of removal of conservative changes suggest that the conservative changes observed in this dataset are evolving via neutral processes in both sexual and asexual lineages. Only radical amino acid changes exhibit severe enough effects on fitness to be differentially maintained in asexual *vs*. sexual lineages, highlighting the utility of distinguishing between mutations with different effects on fitness when comparing the efficacy of selection across lineages.

The observation that asexual *P. antipodarum* accumulate radical nonsynonymous mutations more rapidly than sexual *P. antipodarum* raises the intriguing possibility that asexual *P. antipodarum* might exhibit decreased mitochondrial function compared to sexual counterparts. The presumed severity of some mutations in these genomes (*e.g*., a nonsense mutation in *nad2* of one asexual lineage that would truncate the NAD2 protein by three amino acids) suggests that either mitochondrial function is decreased in at least some of these asexual lineages or that asexuals possess one or more mechanisms to compensate for deleterious mutation load (*e.g*., RNA editing). RNA editing of mitochondrially encoded transcripts has been observed in a variety of plant taxa (Covello and Gray 1989; Gualberto et al. 1989) and in the mitochondrially encoded tRNAs of land snails (Yokobori and Paabo 1995); whether *P. antipodarum* employs similar strategies has not been addressed. Asexual lineages of *P. antipodarum* do exhibit substantial phenotypic variation for mitochondrial function, which has in turn been linked to mitonuclear genotype (Sharbrough et al. 2017). Future work evaluating mitochondrial function at the organelle and organismal levels in *P. antipodarum* will be essential to understanding how the efficacy of selection influences the relative success of sexual *vs*. asexual lineages and the maintenance and distribution of sexual *P. antipodarum*.

### Mitonuclear coevolution in the absence of sex

We interpret these results as a consequence of reduced efficacy of selection in asexual lineages; however, another possible (and non-mutually exclusive) explanation is that the co-transmission (and thus, effective linkage) between the nuclear and mitochondrial genomes in asexuals has facilitated the persistence and spread of beneficial nonsynonymous mutations via selection imposed by cooperation with nuclear-encoded genes (Blier et al. 2001; Meiklejohn et al. 2007). Because apomictic asexuals co-transmit their nuclear and mitochondrial genomes, mutations in either genome may cause decreases in mitochondrial function. Therefore, long-term co-transmission of the nuclear and mitochondrial genomes may provide a scenario in which asexuals experience relatively strong selection favoring compensatory mutation(s). This expectation aligns well with theory suggesting that recombination can be selected against in a scenario where the genes contributing to fitness interact non-additively (see *e.g*., Maynard Smith 1978; Kondrashov 1994; Neiman and Linksvayer 2006). Epistatic selection in which recombination is disfavored has found support from the multiple examples of relatively rapid responses to selection and high heritability of polygenic traits in asexual taxa (*e.g*., Stratton 1991; Browne 1992; Christensen et al. 1992; Stratton 1992; Weeks and Hoffmann 1998). More recently, Lenski and colleagues have shown extensive and rapid adaptive evolution in asexual bacteria lineages in the course of their long-term experimental evolution studies (Lenski 2017). The strong body of work suggesting that mitonuclear interactions are epistatic in nature (*e.g*., Osada and Akashi 2012; Barreto and Burton 2013; Beck et al. 2015; Havird et al. 2015) indicates that the multi-subunit enzymes of the oxidative phosphorylation pathway might be subject to relatively effective evolution by natural selection in an asexual context.

Whether these epistatic interactions contribute to coevolution between the nuclear and mitochondrial genomes of asexual lineages remains an open question. We have not detected any evidence of positive selection acting in the mitochondrial genome of *P. antipodarum* (*e.g*., codon-by-codon *d_N_*/*d_S_* < 1, Neutrality Index > 1 for all 13 protein-coding genes, sliding window *π_A_*/*π_S_* < 1 at all sites; Sharbrough et al. in preparation), though indirect evidence that a particular mitochondrial haplotype is spreading among asexual lineages hints that selection favoring particular mitochondrial haplotypes or mitonuclear combinations might be involved (Paczesniak et al. 2013). Evaluation of rates and patterns of evolution in the nuclear-encoded mitochondrial genes that make up ≥95% of the genes that influence mitochondrial function (Sardiello et al. 2003), coupled with the functional analyses mentioned above, is thus a critical next step towards determining whether mitonuclear linkage in asexuals is at least in part responsible for elevated retention of apparently harmful mutations in mitochondrial genomes.

Ultimately, inefficient removal of seemingly deleterious mutations in asexual lineages is only important to understanding the maintenance of sex and/or the persistence of asexual lineages if those mutations, in fact, negatively affect fitness. Indeed, consistent with the potential that mitonuclear coevolution might be especially effective in asexuals, we cannot exclude the possibility that radical mitochondrial mutations are not as harmful as they appear from our analyses. In particular, we speculate that the selective regime operating on mitochondrial genes may be affected by asexuality itself (*e.g*., elevated heritability of mitonuclear genotypes, allowing for effective compensation of deleterious mutations), such that apparently deleterious mutations do not, in fact, impart reduced function and fitness. Functional validation of deleterious effects of mutations therefore represents an especially central component of determining the evolutionary implications of accelerated accumulation of radical mutations.

### Mitonuclear LD and the efficacy of purifying selection in mitochondrial genomes

For decades, deleterious mutation accumulation has been thought to be an unavoidable hallmark of mitochondrial genome evolution (Gabriel et al. 1993; Neiman and Taylor 2009). In the last few years, several examples of relatively effective natural selection operating in mitochondrial genomes compared to nuclear genomes (Cooper et al. 2015; Konrad et al. 2017) have called this widely held assumption into question. Indeed, the extent to which mitochondrial genomes actually undergo mutational meltdown has profound implications for a number of downstream evolutionary hypotheses including the compensatory model of mitonuclear coevolution (Rand et al. 2004; Dowling et al. 2008; Niehuis et al. 2008; Osada and Akashi 2012; Havird et al. 2015), the evolution of mitonuclear reproductive incompatibilities that could contribute to speciation (Harrison and Burton 2006; Niehuis et al. 2008; Burton and Barreto 2012; Hill 2016), introgressive mitochondrial replacement (Toews and Brelsford 2012; Sloan et al. 2017), and rates of extinction (Lynch et al. 1993; Havird et al. 2016). If natural selection is indeed effective in mitochondrial genomes (especially purifying selection), selection to alleviate deleterious epistatic interactions between nuclear and mitochondrial genomes may not play as central a role in the generation of eukaryotic diversity as previously thought (Montooth et al. 2010; Adrion et al. 2016).

While the results presented here are restricted to mitochondrial genomes, they do provide empirical evidence regarding the efficacy of selection in mitochondrial genomes. In particular, we show that in low *N_e_* conditions (*i.e*., asexual lineages), elevated selective interference appears to increase rates of deleterious mutation accumulation in mitochondrial genomes. That is, as expected under the Hill-Robertson effect, the higher mitonuclear LD experienced by mitochondrial genomes “trapped” in asexual lineages is associated with mitochondrial mutation accumulation. It follows that mitochondrial genomes that appear to experience relatively effective purifying selection (*e.g*., the mitochondrial genomes in sexual lineages of *P. antipodarum*), there must be some mechanism (likely sexual reproduction) whereby the Hill-Robertson effect is ameliorated. Based on this logic, we predict that species harboring relatively few deleterious polymorphisms in mitochondrial genomes (*e.g*., *Caenorhabditis*, *Drosophila*) will also exhibit low levels of mitonuclear LD compared to those harboring many deleterious polymorphisms in mitochondrial genomes (*e.g*., *Nasonia*, *Potamopyrgus*). Characterizing the genetic mechanisms that promote and restrict mitochondrial mutation accumulation in some lineages but not others therefore represents a critical component of understanding the role that mitonuclear coevolution has played in eukaryotic evolution.

## DATA ARCHIVAL LOCATION

Sequence data – GenBank

Python scripts – https://github.com/jsharbrough

## ACKNOWLEDGEMENTS

We thank Cindy Toll and Gery Hehman for their assistance with DNA sequencing. We also thank Stephen I. Wright and Aneil F. Agrawal for helpful discussions regarding interpretation of intraspecific data. We thank Samuel J. Fahrner for helpful discussions regarding bootstrapping. We thank Jeremiah W. Busch and several anonymous reviewers who saw previous versions of this manuscript for their helpful comments. Some of the data presented herein were obtained at the Flow Cytometry Facility, which is a Carver College of Medicine / Holden Comprehensive Cancer Center core research facility at the University of Iowa. The Facility is funded through user fees and the generous financial support of the Carver College of Medicine, Holden Comprehensive Cancer Center, and Iowa City Veteran’s Administration Medical Center. The National Science Foundation (NSF: MCB – 1122176; DEB – 1310825) and the Iowa Academy of Sciences (ISF #13-10) funded this research.

